# Red-light-catalyzed photolipid switching enables leaflet-selective control of membrane mechanics

**DOI:** 10.64898/2026.09.14.751465

**Authors:** Rohit Yadav, Jürgen Pfeffermann, Peter Pohl

**Affiliations:** Institute of Biophysics, Johannes Kepler University Linz, Linz, Austria

## Abstract

Azobenzene photolipids are versatile actuators of membrane mechanics and protein function, yet long-wavelength control and leaflet-selective perturbations remain difficult to impose. Here, we address both limitations. First, we confirm that red-light excitation of a lipidated Nile Blue derivative (NB-lipid) catalyzes rapid *cis*→*trans* photoisomerization of co-localized azobenzene photolipids in planar bilayers without requiring covalent conjugation. Second, we demonstrate that photocatalysis is predominantly intraleaflet, enabling leaflet-selective actuation by incorporating the membrane-targeted chromophore into only one bilayer leaflet. This enables on-demand and rapid writing of transbilayer mechanical asymmetry, which we infer from lock-in capacitance signatures. Critically, symmetric and asymmetric leaflet mechanical perturbations can be evoked within the same membrane by illuminating with blue or red light, respectively. For electron-rich azobenzenes, the anticipated singlet-state photoredox pathway is expected to limit triplet-associated photodamage. Repeated, reversible modulation of gramicidin A, a mechanosensitive peptide ion channel susceptible to reactive oxygen species, supports the compatibility of photocatalysis with membrane protein function. Our results establish a novel tool for dissecting the role of dynamic leaflet-specific mechanical changes in membrane remodeling and protein function.

## 1 Introduction

Biological membranes are inherently asymmetric, both in the lateral dimension and in the transbilayer distribution of lipids. Apart from roles in signaling, this asymmetry allows cells to tune curvature tendencies, store elastic energy, regulate interfacial electrostatics, and shape the lateral pressure profile that couples to membrane proteins [1, 2]. Asymmetric lipid and ionic conditions can generate substantial membrane curvature and differential leaflet tension [3, 4], highlighting the central role of transbilayer asymmetry in membrane remodeling as well as membrane protein and transport regulation [5, 6].

Consequently, transbilayer asymmetry provides a direct mechanical means to regulate membrane protein function, with mechanosensitive channels being particularly sensitive to such perturbations. For example, asymmetric incorporation of lysophospholipids into a single leaflet generates curvature stress that stabilizes the open state of the mechanosensitive channel of large conductance (MscL) even in the absence of applied tension [7]. This behavior reflects the energetic coupling between membrane proteins and their host bilayer: conformational changes incur a bilayer deformation energy that depends on membrane elastic properties and contributes directly to the conformational equilibrium of membrane proteins [8, 9]. Importantly, membrane-mediated regulation does not depend on lipid asymmetry alone: the intrinsic asymmetry of protein shape can itself induce local membrane deformations that feed back on protein function and lateral mobility [10]. These findings motivate examining how asymmetric membrane perturbations influence membrane protein function.

In practice, generating well-defined transbilayer asymmetry in model membranes is feasible, for example by sequential deposition or folding approaches [5, 11], or by microfluidic droplet-based assembly [12]. However, residual solvents or oil phases associated with droplet-based methods can perturb membrane mechanics and protein function [13]. Achieving controlled and quantifiable asymmetry in vesicular systems, and maintaining it over time, remains more challenging [14]. Current strategies rely on lipid exchange methods, such as cyclodextrin-mediated transfer [15], but these approaches are often invasive, difficult to quantify, or limited in temporal control. Overall, it remains difficult to impose programmable, dynamic, rapid, and reversible leaflet-specific perturbations under well-defined conditions, and even more challenging to switch between symmetric and asymmetric membrane perturbations within the same experiment.

Azobenzene-containing photolipids provide a versatile route to optical control of membrane remodeling: reversible *cis*⟷*trans* (*Z* ⟷*E*) photoisomerization alters photolipid molecular structure and thereby switches bilayer material properties [16–19]. Conveniently, these changes can also be monitored with high temporal resolution by capacitance recordings [20] and have been shown to gate ion channels [16, 21]. However, photolipid-induced perturbations are typically imposed symmetrically across both leaflets and rely on direct UV-A/blue-light excitation, which can be a drawback in biological systems. Direct *trans*→*cis* photoisomerization at longer wavelengths has been achieved with appropriately substituted azobenzenes [22, 23]. In red-shifted photolipids, however, weak absorption in the red spectral region can limit switching rates [23]. Alternatively, Shimomura & Kunitake [24] explored an approach to visible-light (580 nm) isomerization of self-assembling azobenzene amphiphiles sensitized by a cyanine chromophore adhered to the bilayer surface. They tentatively attributed visible-light-driven *cis*→*trans* isomerization in the presence of the chromophore to Dexter electron-exchange from the dye’s excited triplet state to interfacial azobenzenes. More recently, Baumgartner *et al.* [25] demonstrated reversible 630 nm-driven *cis*→*trans* switching of AzoPC vesicles doped with 1 mol% C_11_-alkylated Nile Blue.

Here, we extend established non-covalent chromophore-catalyzed bilayer photoisomerization [24, 25] to leaflet-selective membrane switching. We implement this mechanism using NB-lipid, a charged lipidated (C_11_-alkylated) Nile Blue derivative that inserts into membranes and can, in principle, be supplied from the aqueous phase [26]. NB-lipid itself exhibits only very weak protonophoric activity; strong proton transport was observed in combination with the diacylglycerol photolipid OptoDArG, consistent with OptoDArG-assisted transbilayer shuttling of NB-lipid [26]. This enhancement is absent with the phospholipid photolipid OxyAzoPC used in the present experiments. Here we show that under 640 nm illumination, NB-lipid catalyzes *cis*→*trans* isomerization of co-localized azobenzene photolipids without covalent attachment. In contrast to previous examples of catalyzed azobenzene switching that rely on triplet excited states [24, 27–29], NB-mediated switching favors a singlet-state photoredox pathway for redox-matched azobenzenes [30], which we assume also underlies NB-lipid-catalyzed OxyAzoPC switching. Using fast capacitance measurements that primarily report on changes in bilayer area and thickness, we show that efficient photocatalysis requires same-leaflet co-localization of NB-lipid and photolipid, thereby conferring leaflet addressability: selective incorporation of NB-lipid into one leaflet programs red-light responsiveness predominantly in the NB-lipid-containing leaflet, which is bound to generate transbilayer mechanical asymmetry. Following a UV-A reset to the *cis*-enriched state, blue light switches photolipids in both leaflets, allowing reversible transitions between symmetric and asymmetric membrane perturbations on millisecond timescales. Both modes of photoswitching are sufficiently fast to elicit optocapacitive currents driven by rapid changes in bilayer capacitance. Finally, we demonstrate reversible modulation of the mechanosensitive ion channel gramicidin A through repeated whole-bilayer (symmetric) and leaflet-selective (asymmetric) switching. Together, these results establish a general strategy for dynamically controlling leaflet-specific membrane mechanics and probing their role in membrane protein function.

## 2 Results and Discussion

### 2.1 Leaflet-resolved switching in deliberately asymmetric planar lipid bilayers

To establish a baseline for leaflet-resolved perturbations, we constructed deliberately symmetric [16, 32, 33] and asymmetric folded planar lipid bilayers (PLBs) [5, 11, 34] in which we supplied the headgroup-containing azobenzene photolipid OxyAzoPC (chemical structure in Figure 1a) from either only one or both sides during bilayer formation. The main structural lipid was *E. coli* polar lipid extract (PLE), chosen as a robust natural phospholipid mixture that reproducibly forms stable PLBs and provides a chemically heterogeneous membrane matrix for studying photolipid-mediated mechanical perturbations. The preparation of symmetric and asymmetric PLBs is detailed in the Materials and Methods section. The transbilayer redistribution of OxyAzoPC is negligible over the timescale of our experiments since the spontaneous flip-flop of the phosphatidylcholine lipids is generally slower than the lateral exchange between the individual membrane leaflets and the connected lipid monolayer on the PTFE support [11, 35]. More specifically, for the closely related headgroup-containing azobenzene phospholipid azo-PC, PMF calculations yielded translocation barriers of 47–50 kJ mol^−1^ for both *cis* and *trans* states, corresponding to flip-flop on the hour timescale [19]. These times are several orders of magnitude longer than the millisecond-to-second switching times analyzed here. In addition, we previously used OxyAzoPC in asymmetric planar bilayers and observed a stable transmembrane boundary-potential difference over time [26]; in the same study, OxyAzoPC did not support the pronounced NB-lipid-mediated proton conductance observed with the rapidly flip-flopping photolipid OptoDArG. Hence, OxyAzoPC was expected to remain enriched in the leaflet at the side(s) it was supplied to on the experimental timescale. We probed the membrane structural response to photolipid switching indirectly by high-frequency lock-in measurements of membrane capacitance *C* (a schematic of the experimental setup is shown in Figure 1b). Photoisomerization changes the molecular geometry and preferred area of the photolipids and thereby alters bilayer area and thickness, which contribute to changes in *C* [16, 20]. Capacitance reports the combined effects of bilayer area *A*, thickness *d*, and permittivity *ϵ* through *C* = *ϵA*/*d*. We interpret the photoinduced response primarily in terms of changes in *A* and *d*, consistent with previous structural and electrical measurements [16, 20, 26].

**Figure 1:**
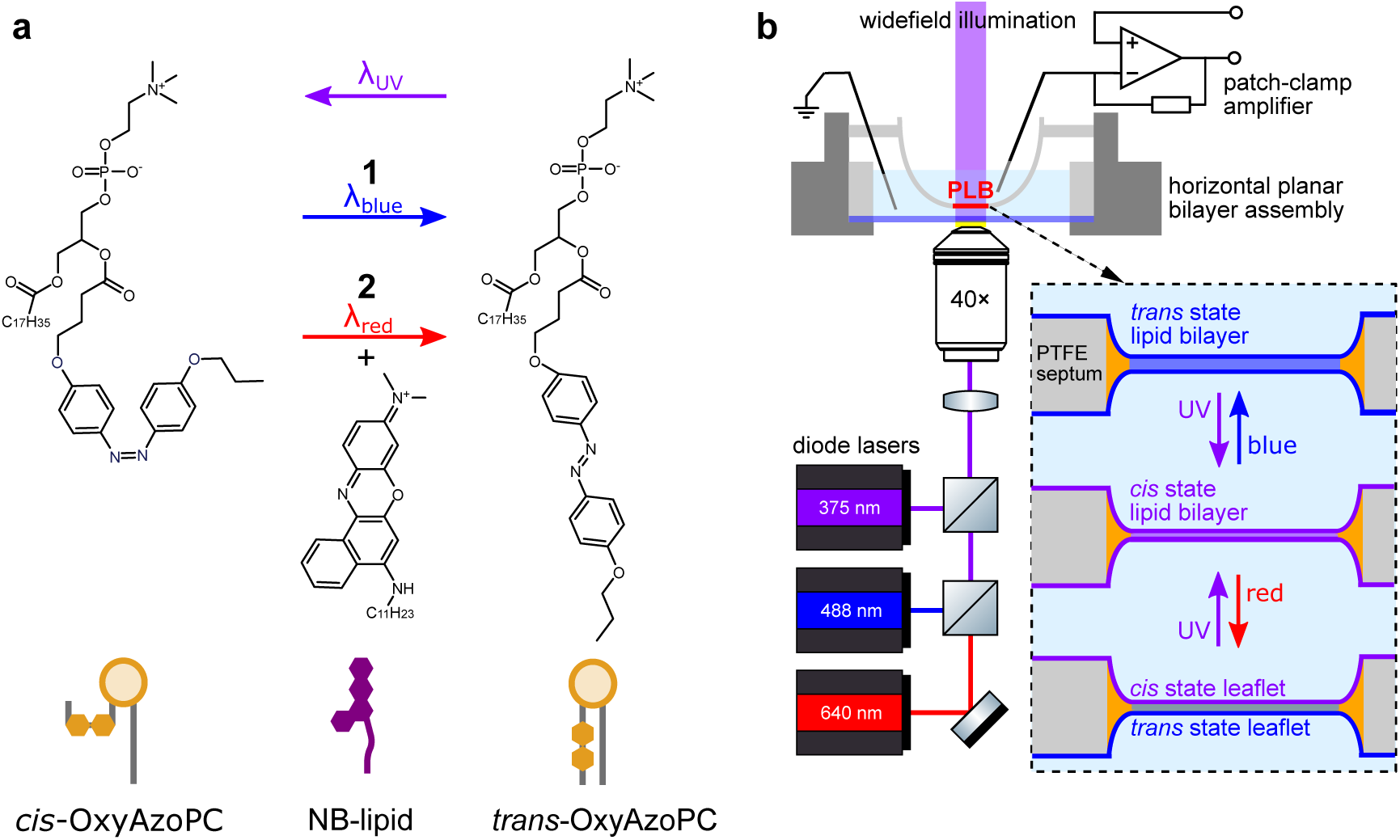
**a,** *cis*-OxyAzoPC is generated from *trans*-OxyAzoPC by UV-A light exposure. To isomerize from *cis* to *trans*-OxyAzoPC, two avenues are possible: **1,** direct blue-light excitation; **2,** red-light-catalyzed isomerization by NB-lipid (center molecule). Schematics representing the photolipid states and photocatalyst are shown below the structures. **b,** Schematic of the experimental setup used for rapid photoisomerization of photolipid-containing planar lipid bilayers and simultaneous current measurements. For UV-A and blue-light exposure, ‘*cis*’ and ‘*trans*’ refer to the *cis*-enriched and *trans*-enriched photostationary states achieved with 375 nm and 488 nm light, respectively [31].

First, in symmetric bilayers with 10 wt% OxyAzoPC in both leaflets, blue (488 nm) illumination produced a reproducible decrease in *C* consistent with *cis*→*trans* photoisomerization of the photolipid (Figure 2a) [16, 20]. The fractional decrease in capacitance was around 1.7%, and because we used high-irradiance illumination, *C* changed on the millisecond timescale. In asymmetric bilayers with 10 wt% OxyAzoPC restricted to one leaflet, blue light likewise produced a decrease in *C* but with a smaller amplitude of 0.9% (Figure 2a). Thus, restricting direct photolipid excitation to one leaflet produces roughly half the capacitance change of symmetric bilayer switching. As anticipated, UV-A (375 nm; 50 ms pulses of UV-A light were used throughout this manuscript to establish *cis*-state PLBs) illumination reverses blue-light-evoked changes in *C* (Figure S1), confirming reversible *cis*⟷*trans* photochemistry. This response was independent of membrane orientation: comparable fractional capacitance changes and relaxation kinetics were obtained when OxyAzoPC was confined to either the upper or lower leaflet (Figure S1). Together, these results show that confining OxyAzoPC to one leaflet preserves rapid, reversible switching while reducing the whole-bilayer capacitance response, providing a baseline for evaluating leaflet-specific photocatalysis.

**Figure 2:**
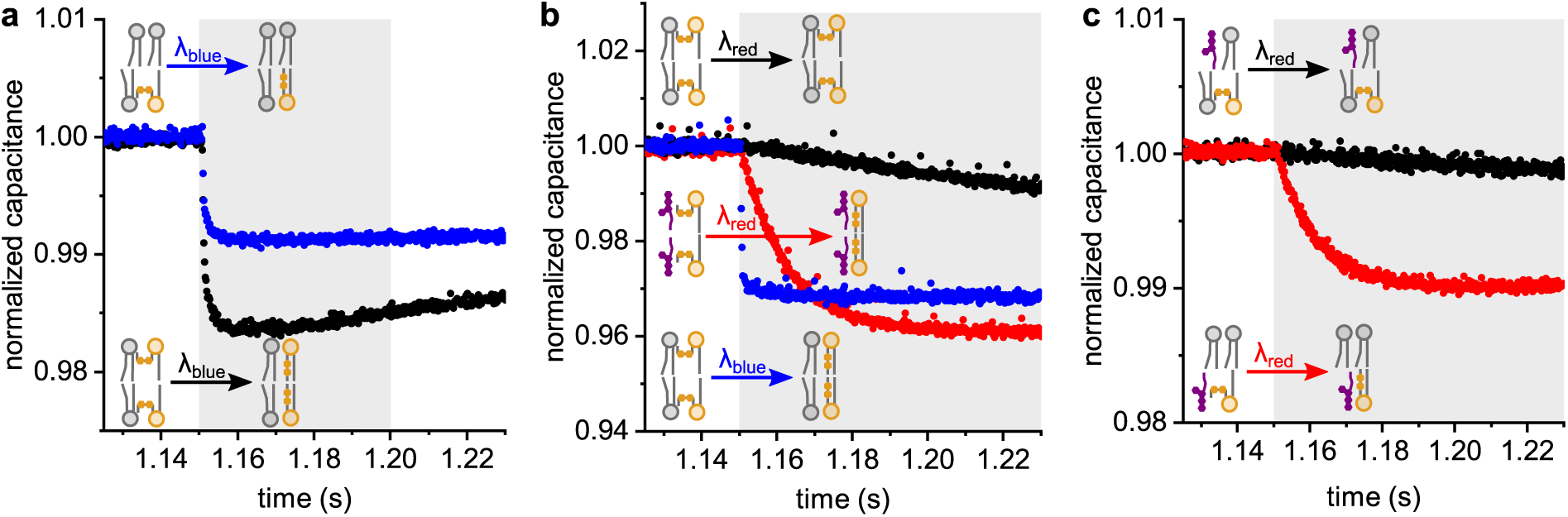
NB-lipid catalyzes leaflet-specific red-light *cis*→*trans* switching of azobenzene photolipids in lipid bilayers. **a,** Lock-in capacitance recordings of *E. coli* PLE bilayers with 10 wt% OxyAzoPC in one (blue trace) or both (black trace) leaflets. Prior to blue-light exposure, the bilayer was switched to the *cis* state with a 50 ms UV-A light pulse (not shown). A 50 ms 488 nm pulse starting at 1.15 s decreases *C* due to photolipid photoisomerization. The smaller amplitude change in the blue trace reflects restriction of the photolipid to one leaflet. The lower-capacitance plateau reached after blue-light exposure corresponds to the *trans*-enriched photostationary state, whereas UV-A illumination restores the higher-capacitance *cis*-enriched state (not shown). **b,** Symmetric bilayers with 20 wt% *cis*-OxyAzoPC per leaflet, 50 ms blue/1 s red light pulse starting at 1.15 s: red light yields a minimal reduction in *C* (black trace), whereas blue light switches rapidly (blue trace). With 1 wt% NB-lipid in both leaflets, red light drives fast photocatalyzed switching with a larger capacitance decrease than direct blue-light excitation (red trace), consistent with the near-quantitative conversion reported for the corresponding molecular system [30]. **c,** Efficient photocatalysis requires leaflet co-localization: red-light switching (1 s pulse starting at 1.15 s) is slow when *cis*-OxyAzoPC (10 wt%) and NB-lipid (1 wt%) are in opposite leaflets (black) and fast when both are in the same leaflet (red), indicating weak catalysis across the bilayer midplane.

### 2.2 NB-lipid catalyzes leaflet-specific photolipid switching

As anticipated, in *cis*-OxyAzoPC-containing bilayers, red (640 nm) light illumination resulted in very slow capacitance switching (black trace in Figure 2b), consistent with inefficient direct excitation of *cis*-azobenzene. By contrast, blue illumination of the same membrane drove rapid changes in *C* owing to efficient direct excitation (blue trace in Figure 2b). Consistent with the vesicle-scaffolded sensitization reported by Baumgartner *et al.* [25], red-light responsiveness increased markedly when the lipidated Nile Blue (NB) derivative NB-lipid (structure in Figure 1a) was co-confined with the photolipids. Lipidation of the phenoxazine dye NB increases NB-lipid’s hydrophobicity and favors partitioning into lipid bilayers [26], where NB-lipid is in close proximity to photolipid azobenzenes. The singlet excited-state reduction potential reported for NB (*E*^S1^_red_ = 1.10 V versus ferrocene) slightly exceeds the oxidation potential of a representative *para,para*-bisalkoxy azobenzene (*E*^S0^_ox_ = 1.07 V versus ferrocene), which has the same electron-rich substitution pattern as the azobenzene in OxyAzoPC [30]. This redox compatibility, together with NB’s low intersystem crossing quantum yield (Φ_ISC_ < 0.03 [36]), supports singlet-state photoredox as a plausible pathway for NB-lipid-catalyzed OxyAzoPC switching [30]. In this proposed route, singlet-excited NB accepts an electron from the azobenzene, allowing radicalcation isomerization followed by back electron transfer. The proposed singlet-photoredox route differs from the triplet-mediated photoredox and energy-transfer mechanisms reported for other sensitized azobenzene systems [24, 27, 37]. Critically, at 1 wt% NB-lipid in the lipid mixture used for planar bilayer formation, red light caused a fast reduction in *C* (red trace in Figure 2b), approaching the rate achieved by direct blue-light excitation. This response is consistent with rapid photolipid *cis*→*trans* isomerization driven by co-confined NB-lipid photocatalysts.

Azobenzene photolipids do exhibit thermal *cis*→*trans* isomerization, but this process is too slow [38] to interfere with our measurements of changes in *C* occurring on the millisecond timescale. Yet, excitation of NB-lipid can cause modest membrane heating and could accelerate thermal isomerization. To quantify this contribution, we exposed PLBs of the following nominal compositions to 1 s of red laser light: (i) PLE with 1 wt% NB-lipid and no photolipid (Figure S2a), as well as (ii) PLE with neither NB-lipid nor photolipid (Figure S2b). In the NB-lipid-containing PLB, *C increased* by approximately 0.35% within the first 80 ms. This corresponds to an estimated temperature increase of approximately 1 ^∘^C based on the reported thermal capacitance increase rate of ∼0.3 % ^∘^C^−1^ observed across different membrane systems [39]. Importantly, this thermal response is much smaller than, and opposite in sign to, the ∼4% capacitance *decrease* observed in NB-lipid/OxyAzoPC membranes under red illumination. Photothermal heating therefore cannot account for the observed switching response. For a lipid bearing two azobenzene-containing chains in liposomes, Moss & Jiang [38] report a 2.4-fold increase in the thermal back-isomerization rate upon heating from 21 ^∘^C to 30 ^∘^C, yet the half-life remains approximately 2 h at 30 ^∘^C. Such a modest acceleration of a slow process, particularly with the estimated heating here of only about 1 ^∘^C, cannot explain the millisecond capacitance response; light-driven photoisomerization can.

Notably, a proximity-dependent photoredox mechanism in our system can be readily rationalized: at 1 wt% NB-lipid, corresponding to ∼1.5 mol% in the bacterial lipid extract used, about 9% of photolipids are expected to have an NB-lipid as a direct nearest neighbor at any given instant, assuming random mixing and about six nearest neighbors per lipid. However, the mean lateral spacing between NB-lipids is only ∼8 nm. Assuming lateral diffusion coefficients of order 10 *μ*m^2^ s^−1^ for both species, the characteristic NB-lipid/photolipid encounter time is below 1 µs. Thus, during the red-light illumination period (50 ms to 1 s were used in this study), photolipids repeatedly encounter NB-lipids, providing a plausible basis for efficient intermolecular photocatalysis. NB-lipid thus enables efficient red-light switching of OxyAzoPC, with the favored singlet-state pathway expected to limit triplet-associated photodamage [25, 40, 41].

Photocatalysis was also sufficiently efficient to generate optocapacitive currents arising from rapid changes in membrane capacitance [20]. Symmetric bilayers containing *cis*-OxyAzoPC and NB-lipid exhibited robust optocapacitive currents under direct blue-light excitation and clear, albeit smaller, currents under red-light illumination (Figure S3), reflecting slower NB-lipid-catalyzed switching (*τ* ∼ 5 ms with red light versus *τ* = 0.24 ms with blue light). Further, red-light switching produced larger fractional capacitance decreases than direct blue excitation (Figure 2b), consistent with a higher achieved *trans*-OxyAzoPC content under photocatalysis. Near-quantitative *cis*→*trans* conversion of *para,para*-bisalkoxy azobenzenes by NB was reported for the corresponding molecular system, whereas the photostationary state of the same azobenzene under 490 nm excitation contained only ≈82% *trans* [30].

Notably, photocatalysis was not restricted to *para,para*-bisalkoxy azobenzenes and OxyAzoPC. We also observed red-light photocatalyzed switching in bilayers containing 20 wt% OptoDArG and 1 wt% NB-lipid in both leaflets; OptoDArG contains two *para,para*-dialkyl azobenzenes (molecular structure and capacitance record in Figure S4) [16, 20, 21, 26, 42]. This is consistent with reported red-light photocatalysis in AzoPC vesicles doped with 1 mol% NB-lipid, since AzoPC shares the same *para,para*-dialkyl azobenzene substitution pattern as OptoDArG [25]. However, the reported oxidation potentials of *para,para*-dialkyl azobenzenes make their photooxidation by singlet-excited NB unfavorable [30]. Hence, their catalyzed isomerization is not expected to proceed via singlet-state photoredox. Instead, Baumgartner *et al.* [25] proposed that singlet-excited NB forms a transient excited complex (exciplex) with the azobenzene. In their proposed mechanism, intersystem crossing within this complex produces a triplet exciplex that subsequently populates the azobenzene triplet, whose rapid relaxation favors the *trans* state. This singlet-initiated, triplet-mediated pathway does not require a long-lived NB triplet, consistent with the photostability reported for NB-lipid/AzoPC vesicles [25]. Together, these observations support singlet-excited NB-lipid-catalyzed switching of both bisalkoxy- and dialkyl-substituted azobenzene photolipids, although the microscopic pathways remain tentative.

Returning to OxyAzoPC, we tested whether efficient photocatalysis requires the photolipid and NB-lipid in the same leaflet. Therefore, we prepared asymmetric planar lipid bilayers by folding two separately prepared monolayers [5, 11, 34]. For the same-leaflet configuration, both compounds were included in the lipid mixture used to form one monolayer. For the opposite-leaflet configuration, OxyAzoPC was included in the mixture for one monolayer and NB-lipid in the mixture for the other. We found that photocatalysis was reduced by approximately one order of magnitude when NB-lipid and OxyAzoPC were placed in opposite leaflets compared to the same-leaflet configuration, as reflected by the markedly slower rate of photoisomerization (Figure 2c). Approximating the capacitance decay within the first 500 ms of red-light exposure by a single exponential, we determined an apparent capacitance-response time constant *τ* = 18 ± 7 ms for leaflet co-localization and *τ* = 181 ± 42 ms for OxyAzoPC and NB-lipid in opposite leaflets (both numbers are mean±SD of three records from three independent preparations). The latter *τ* is only modestly shorter than that in OxyAzoPC-containing bilayers without any NB-lipid at all (see black trace in Figure 2b), for which we obtained *τ* = 255 ± 34 ms (mean±SD of four records from four independent preparations). This strong leaflet dependence is consistent with predominantly intraleaflet photocatalysis and reduced catalytic efficiency across the hydrophobic membrane midplane. This observation supports a mechanism requiring close molecular contact. Taking into consideration the redox compatibility discussed above, photoinduced electron transfer between neighboring molecules during the nanosecond singlet excited-state lifetime of NB emerges as a prime candidate [30]. These interactions are likely facilitated by two-dimensional co-confinement within the membrane plane, where sub-µs NB-lipid–photolipid encounter times permit repeated transient interactions during red-light illumination.

### 2.3 Wavelength-selective photolipid switching controls transbilayer mechanical asymmetry

Having established predominantly intraleaflet photocatalysis, we next tested whether asymmetric incorporation of NB-lipid would allow illumination wavelength to select between symmetric and asymmetric membrane mechanical actuation. Therefore, we compared capacitance signatures under blue and red light in bilayers containing 20 wt% OxyAzoPC in both leaflets but 1 wt% NB-lipid in only one leaflet. In these membranes, blue light produced a rapid decrease in *C* of amplitude 4.3 ± 0.1% (mean±SD of five records from two independent preparations), consistent with symmetric switching, whereas red light resulted in a slower decrease and a lower fractional reduction in *C* of 3.3 ± 0.1% (Figure 3a).

**Figure 3:**
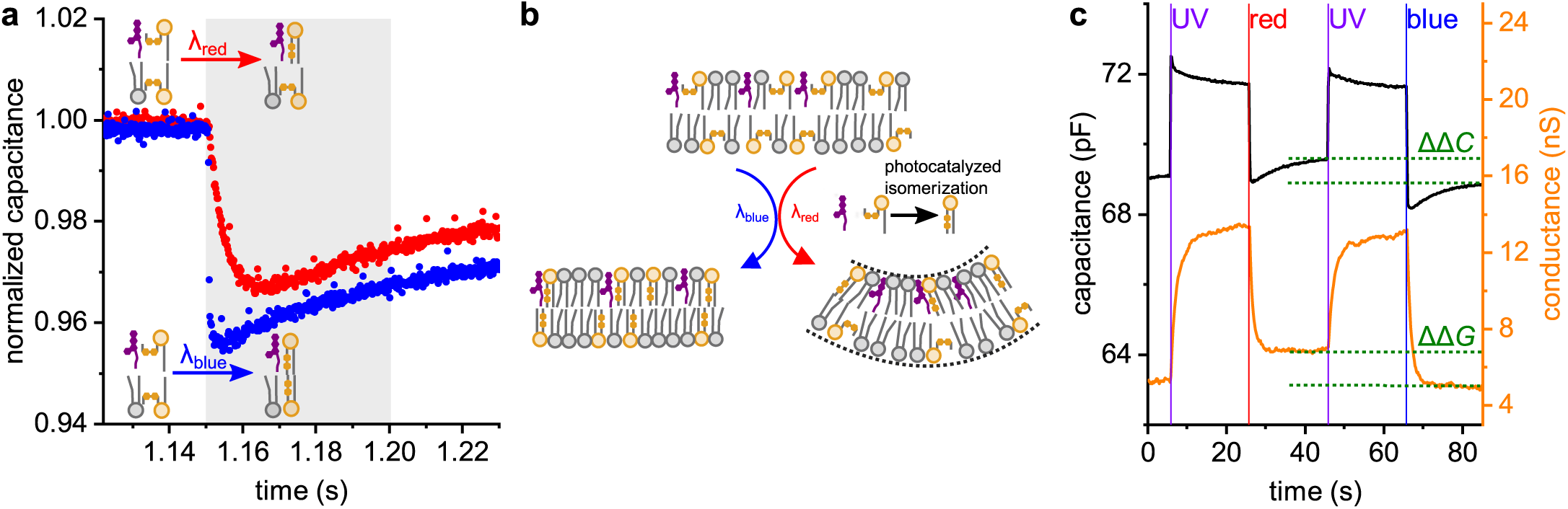
Wavelength-dependent on-demand symmetric or asymmetric bilayer photoisomerization. **a,** In asymmetric bilayers with 20 wt% OxyAzoPC per leaflet and 1 wt% NB-lipid in only one leaflet, blue light (50 ms pulse starting at 1.15 s) produces a rapid decrease in *C* consistent with symmetric switching (blue points), whereas red light (1 s pulse starting at 1.15 s) results in a slower decrease and a lower fractional decrease in *C* (red points). This is consistent with leaflet-selective NB-lipid-catalyzed switching. **b,** Photolipid photoisomerization enables symmetric actuation by direct blue-light excitation, whereas NB-lipid-mediated photocatalysis under red light predominantly switches the NB-lipid-containing leaflet (asymmetric actuation). The expected reduction in preferred area of the switched leaflet favors bending toward that leaflet, placing it on the concave side. The curved configuration illustrates this expected mechanical tendency or leaflet stress asymmetry; macroscopic membrane deformation was not observed in our experiments. The actual deformation depends on membrane tension and boundary constraints; tension preferentially suppresses long-wavelength undulations without eliminating shorter-wavelength modes [43]. **c,** The incorporation of gramicidin A ion channels increases membrane conductance *G*. The bilayer was isomerized with 50 ms pulses of UV-A (*trans*→*cis*), red, or blue light (*cis*→*trans*). The graph shows an average of six superimposed UV-A/red/UV-A/blue switching cycles (the full record is shown in Figure S5). The delayed decrease in *G* is consistent with reduced gramicidin A activity as the membrane thickens. The effect of photocatalyzed asymmetric switching on *G* and *C* is reduced relative to symmetric actuation by blue light.

As apparent in Figure 3a, the reduction in *C* under red light is greater than 50% of that under blue light. Three factors may contribute to this deviation from the simplest one-half expectation: (i) Red-light photocatalysis is expected to drive close-to-quantitative *cis*→*trans* switching, whereas direct blue-light excitation produces a photostationary state with a lower *trans* content [30]. (ii) Red light also induces weak direct photoisomerization of *cis*-OxyAzoPC in the nominally NB-lipid-free leaflet (cf. Figure 2b). (iii) The two leaflets are elastically coupled and therefore do not respond independently [34, 44]. Photolipid photoisomerization changes molecular geometry, preferred molecular area, thickness, and lateral packing [16, 19]. Consequently, leaflet-selective switching can induce an elastic readjustment of the opposing leaflet, and the whole-bilayer capacitance response to asymmetric actuation need not be exactly one half of that obtained upon symmetric switching.

The capacitor relation (*C* = *ϵA*/*d*) helps specifying the mechanical interpretation. If changes in *ϵ* are small [26], the observed decrease in *C* implies a decrease in *A*/*d*. Previous work on OptoDArG and related azobenzene photolipids associates *cis*→*trans* switching with thickening and a reduction in preferred area per lipid [16, 19]. Leaflet-selective switching of OxyAzoPC is therefore expected to reduce the preferred area of the switched leaflet relative to the opposing leaflet. In the bilayer-couple picture, this area mismatch favors bending toward the switched leaflet, placing it on the concave side and the leaflet of larger preferred area on the convex side (Figure 3b) [45]. Macroscopic bending may occur in an unconstrained, tension-free membrane; the extent and spatial form of deformation depend on membrane tension, bending rigidity, and boundary constraints, with tension preferentially suppressing long-wavelength undulations [43]. Whilst we did not directly observe macroscopic membrane deformation in our experiments, Figure 3b illustrates a rational expected mechanical tendency towards bending arising from leaflet-selective photolipid switching.

### 2.4 Wavelength-selective membrane actuation differentially affects a molecular force probe

Finally, we compared the effects of symmetric (blue) and asymmetric (red) actuation on the mechanosensitive peptide ion channel gramicidin A (gA) [16]. Because gA is sensitive to reactive oxygen species (ROS)-mediated photoinactivation [40, 46], these measurements also test whether channel function is retained during repeated photocatalyzed switching. Therefore, we prepared planar bilayers containing 20 wt% OxyAzoPC in both leaflets and 1 wt% NB-lipid in only one leaflet, and added 0.1 nM gA to the aqueous compartments on both sides of the membrane from a 100 nM stock solution in ethanol (Figure 3c). Gramicidin A spontaneously partitions from the aqueous phase into the bilayer leaflets and forms cation-conducting channels by transbilayer dimerization. Simultaneous lock-in capacitance *C* and conductance *G* recordings evidenced (i) a rapid decrease in *C* upon blue- or red-light exposure that is reversible by UV-A light, and (ii) a delayed decrease in *G*, consistent with slower re-equilibration of gA monomers and conducting dimers [47]. The decrease in *C* is qualitatively consistent with membrane thickening, which increases hydrophobic mismatch and disfavors conducting gA dimers, in agreement with an earlier study [16]. Besides hydrophobic mismatch, photolipid isomerization affects bending rigidity and spontaneous curvature [16], which is anticipated to affect curvature elastic stress and thereby the membrane deformation energy governing gA channel formation and lifetime [16, 47–49]. Importantly, (i) the reduction in both *C* and *G* was larger under blue light than under red light, consistent with symmetric versus asymmetric actuation, and (ii) no cumulative loss of gA activity was detected, consistent with limited photodamage and ROS generation (Figure S5 shows the full electrical record). This is qualitatively consistent with the reported photostability of NB-lipid in AzoPC vesicles under prolonged red-light exposure [25]. Also, note that the distinct red- and blue-light responses persisted throughout repeated switching cycles (Figure 3c and Figure S5), consistent with retained leaflet selectivity. In conclusion, the same bilayer can be actuated in a wavelength-selective manner to produce symmetric or asymmetric mechanical perturbations that can differentially modulate membrane protein activity.

## 3 Conclusion

We demonstrated that conjugation-free red-light driven photocatalysis by 2D membrane confinement of photolipids and photocatalysts can be restricted to one bilayer leaflet, enabling leaflet addressability: Selective incorporation of NB-lipid into one bilayer leaflet allows red illumination to drive switching mainly in the NB-lipid-containing leaflet. Exposing the same bilayer to blue light switches photolipids in both leaflets symmetrically. Illumination wavelength thus becomes a flexible selector for mechanical actuation: blue light produces symmetric membrane mechanical changes via direct photolipid excitation in both leaflets, whereas red light switches photolipids predominantly in the NB-lipid-containing leaflet and is thus expected to generate leaflet-selective changes in preferred molecular area and asymmetric mechanical stresses within the bilayer leaflets. Capacitance measurements readily distinguish these symmetric versus asymmetric perturbations and report switching kinetics with millisecond temporal resolution.

We believe that this wavelength-selective control of symmetric and asymmetric photoisomerization provides a versatile tool for dissecting how leaflet-specific mechanical changes regulate membrane remodeling and membrane protein function. The ability to dynamically write transbilayer asymmetry on demand within the same membrane, coupled with rapid electrical readouts, enables previously inaccessible tests and direct comparison of how membrane proteins such as mechanosensitive channels respond to symmetric versus asymmetric leaflet perturbations. Future explorations of non-flip-flopping photocatalysts could extend these capabilities to liposomes and living cells by enabling post-formation imposition of on-demand leaflet addressability.

## 4 Materials and Methods

### 4.1 Compounds

*E. coli* Polar Lipid Extract (PLE, item no. 100600) was purchased from Avanti Polar Lipids (distributed by Merck). OxyAzoPC (1-stearoyl-2-[4-(4-((E)-(4-propoxyphenyl)diazenyl)phenoxy)-butanoyl]-*sn*-glycero-3-phosphocholine; see Figure 1a; previously used in [26]) and NB-lipid (compound **S43** in [25]; synthesis and characterization described therein) were provided by the group of Dr. Oliver Thorn-Seshold. OptoDArG [16, 20, 21, 26, 42] was synthesized and provided by the laboratory of Dr. Toma Glasnov. Lipid stock solutions and mixtures were aliquoted and mixed in chloroform in amber glass microreaction vials. Prior to storage at −80 ^∘^C, solvent was removed under a mild vacuum gradient (Rotavapor, Büchi Labortechnik AG) and the residual dried lipids were flooded with argon. A 100 nM solution of gramicidin A from *Brevibacillus brevis* (SKU 50845, Fluka) in ethanol was kept at −20 ^∘^C. Aqueous buffers (150 mM KCl, 10 mM HEPES pH 7.4) were prepared fresh from laboratory-grade dry substances (VWR, Merck, or Fisher Scientific) dissolved in ultrapure water (>18 M*Ω* cm, Milli-Q water purification system) and pH-adjusted (FiveEasy, Mettler Toledo).

### 4.2 Horizontal planar lipid bilayer setup

Solvent-depleted horizontal PLBs (specific capacitance >0.75 µF cm^−2^) were formed by the Montal–Mueller folding technique [20, 32, 34] in a custom PTFE chamber. Figure S6 shows a schematic of the horizontal PLB formation process. The upper and lower compartments contained 380 and 1500 µL of aqueous buffer, respectively, and were separated by a 25 µm-thick PTFE diaphragm containing an approximately 70 µm-diameter aperture created by high-voltage discharge. Before bilayer formation, the diaphragm was treated with 0.6 vol% hexadecane in hexane. Hexane was allowed to evaporate for at least 30 min, leaving a residual hexadecane solvent annulus around the aperture that anchors the bilayer within the aperture [50]. Thus, the membranes are solvent-depleted rather than strictly solvent-free. The treated septum was fixed with silicone paste to the underside of the upper compartment.

Lipid mixtures in hexane (10 mg mL^−1^) were applied onto the air–water interfaces of the two compartments. For asymmetric PLBs [5, 11, 34], different lipid mixtures were applied to the aqueous interfaces of the lower and upper compartments. The main structural lipid was *E. coli* PLE. In experiments on PLBs containing OxyAzoPC, OptoDArG and/or 1 wt% NB-lipid, these compounds were added to the lipid mixtures spread at the interfaces, never to the aqueous phases. OxyAzoPC and OptoDArG were present in their stable *trans* state during lipid mixture preparation and the application to the aqueous interfaces. NB-lipid is a Nile Blue derivative and does not undergo *cis*–*trans* photoisomerization.

After hexane evaporation (>30 min), each aqueous surface was covered by its separately prepared lipid monolayer. The PTFE diaphragm supporting the upper aqueous compartment was then rotated such that the two monolayers were brought into apposition at the aperture. Their hydrophobic faces thereby contacted each other and formed the planar bilayer, while the surrounding hexadecane annulus remained at the aperture rim; the folding of horizontal PLBs is schematically illustrated in Figure S6. Each leaflet remains continuous with lipid monolayers on top of the torus and PTFE septum; lipids are diffusively exchanged between a PLB leaflet and its adjoining monolayer. When different lipid mixtures were applied to the aqueous interfaces, those monolayers had different compositions, which in turn controlled the composition of the connected bilayer leaflets. The strong acceleration of photoisomerization when NB-lipid and OxyAzoPC are supplied to the same interface rather than opposite interfaces supports substantial leaflet asymmetry (Figure 2c), although the precise leaflet compositions were not quantified. Functional incorporation of NB-lipid was confirmed by the strong acceleration of red-light-induced OxyAzoPC photoisomerization. All reported lipid and NB-lipid concentrations are nominal values for the mixtures used to form each leaflet.

The lower compartment was sealed at the bottom by a 30 mm-diameter cover glass (No. 1, Assistent, Hecht Glaswarenfabrik GmbH & Co KG) secured with a threaded PTFE ring. The chamber was mounted on the stage of an Olympus IX83 inverted microscope equipped with an iXon 897 E EMCCD camera (Andor, Oxford Instruments Group). The PLB was positioned within the working distance of a 40×/0.65 NA infinity-corrected plan achromat air objective (PLN40X, Olympus) by micrometer screws on the chamber holder. Laser triggering and electrophysiological acquisition were synchronized via the motorized microscope’s real-time controller (U-RTC, Olympus) running cellSens software (Olympus). All experiments were performed at room temperature.

### 4.3 Electrical recordings

For electrical recordings, Ag/AgCl agar salt bridges (0.5 M KCl) were placed in each compartment and connected to the headstage of an EPC 9 patch-clamp amplifier (HEKA Elektronik, Harvard Bioscience); headstage and chamber were enclosed in a Faraday cage. Voltage-clamp recordings were acquired with PATCHMASTER 2x91 (HEKA Elektronik, Harvard Bioscience), with currents filtered at 10 kHz (Bessel) and digitized at 25–50 kHz. The HEKA software lock-in was used for capacitance and conductance measurements. Data were exported from PATCHMASTER and subsequently analyzed and plotted in Mathematica 14.3 (Wolfram Research) and OriginPro 2026 (OriginLab Corporation).

### 4.4 Light exposure of PLBs

Three digitally modulated diode lasers (TOPTICA Photonics) were used for PLB illumination: 488 nm (iBEAM-SMART-488-S-HP, blue), 375 nm (iBEAM-SMART-375-S, UV-A), and 640 nm (iBEAM-SMART-640-S, red). All beams were coupled into the back-focal plane of the objective through the ZET488/640rpc main dichroic mirror (Chroma, AHF analysentechnik). Spatial filters were removed from all beam paths to maximize power at the sample stage. Beam radii (at 1/e^2^ intensity) and incident powers at the sample stage were: UV-A, roughly 78 µm at ∼18 mW; blue, roughly 41 µm at ∼69 mW; red, roughly 68 µm at ∼80 mW.

## Author Contributions

RY performed experiments. RY and JP analyzed data. JP wrote the initial draft. PP conceived the project. All authors designed experiments and contributed to writing the manuscript.

## Acknowledgments

We thank Dr. Oliver Thorn-Seshold (TU Dresden) for providing NB-lipid and OxyAzoPC and Dr. Toma Glasnov (University of Graz) for providing OptoDArG. This research was funded in whole by the Austrian Science Fund (FWF) grants P34826 and P36399 to Peter Pohl.

## Data Availability Statement

The data supporting the findings of this study are available from the corresponding author upon reasonable request.

## Supplementary Information

**Figure S1:**
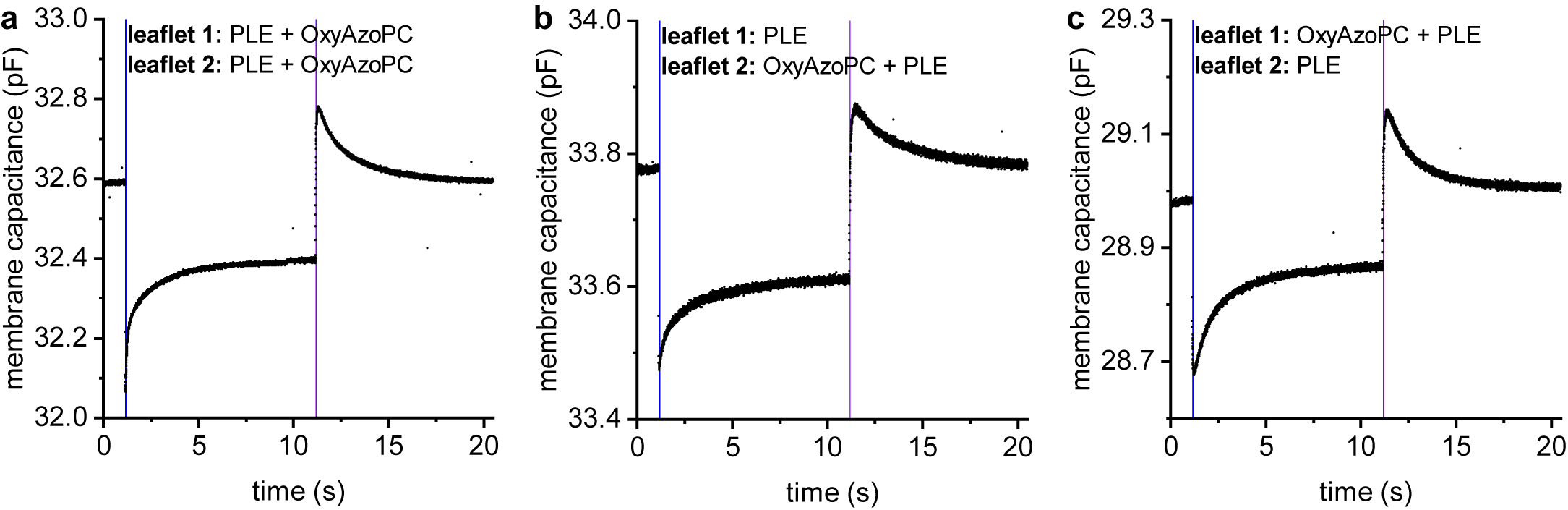
Unnormalized capacitance recordings underlying the leaflet-resolved switching experiments. Raw experimental capacitance recordings (septum capacitance was subtracted; for display, the traces were downsampled by replacing each nonoverlapping block of 10 consecutive data points with its mean) on *E. coli* PLE bilayers containing 10 wt% OxyAzoPC in both (**a**) or only one leaflet (**b** and **c**). Blue and UV-A light pulses lasted 50 ms. **b,** Here, OxyAzoPC was added to the lipid mixture forming the lower leaflet. The blue curve in Figure 2a was prepared from this recording. **c,** Here, OxyAzoPC was added to the lipid mixture forming the upper leaflet. Reversing the orientation of the asymmetric membrane produced comparable fractional capacitance changes and relaxation kinetics upon photolipid switching. The symmetric bilayer produces the larger capacitance change reported in Figure 2a. The slower capacitance relaxation following the initial photoinduced step reflects mechanical readjustment of the bilayer–torus system, as described previously [20, 21].

**Figure S2:**
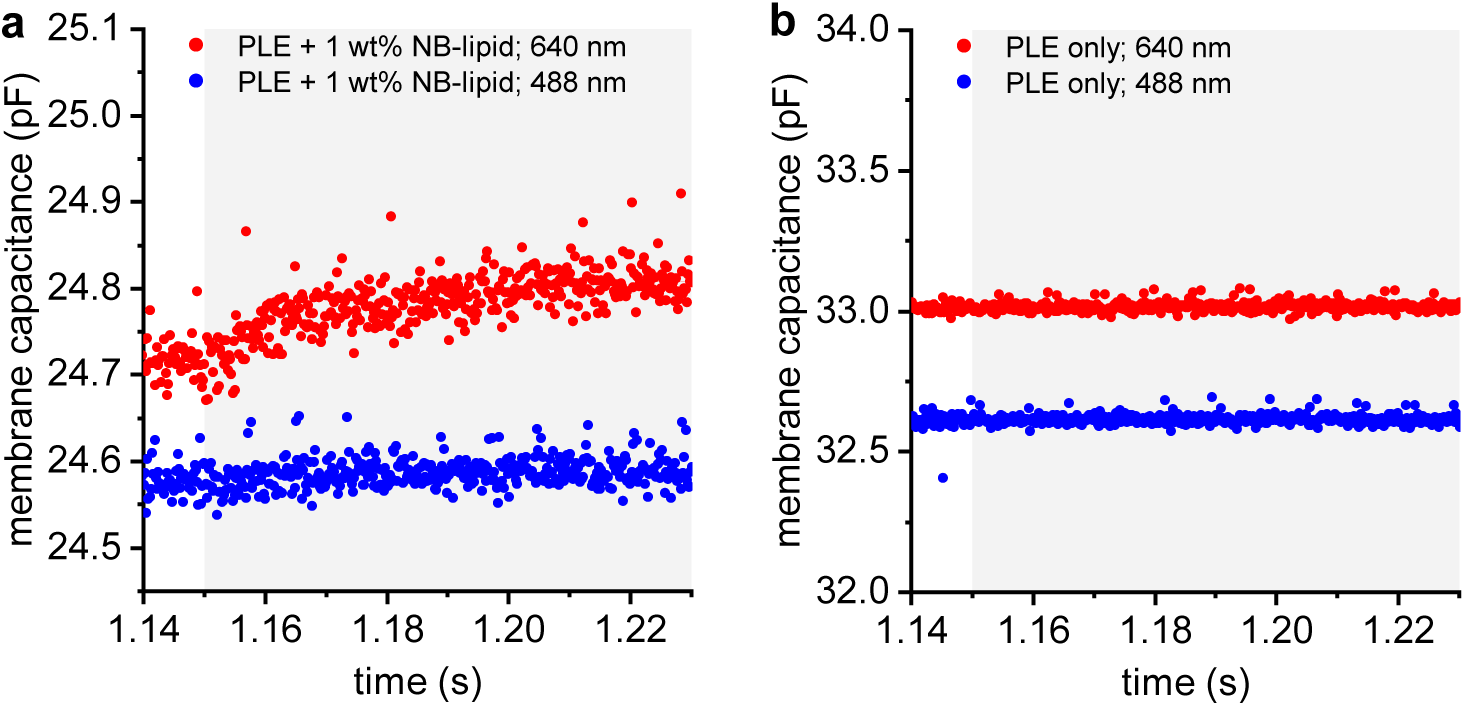
Control capacitance measurements. **a,** Lock-in capacitance recordings of an *E. coli* PLE bilayer with 1 wt% NB-lipid in both leaflets. Starting at 1.15 s, the PLB was exposed to blue (blue trace) or red light (red trace) for 1 s. Upon blue exposure, capacitance remained practically unchanged within the first 80 ms. Upon red excitation, capacitance *increased* by 0.35% within the first 80 ms of exposure, as compared to the ≈4% *decrease* in capacitance in the presence of 20 wt% OxyAzoPC (red trace in Figure 2b). This increase in capacitance with red light indicates a modest increase in temperature due to the excitation of NB-lipid because, under well-defined conditions, changes in membrane capacitance directly report on changes in membrane temperature [51]. Using the reported thermal capacitance coefficient of approximately 0.3 % ^∘^C^−1^ observed across different membrane systems [39], this corresponds to an estimated temperature increase of approximately 1 ^∘^C within the first 80 ms of red-light exposure. **b,** Without NB-lipid (and without OxyAzoPC), there is no capacitance response to either blue or red light.

**Figure S3:**
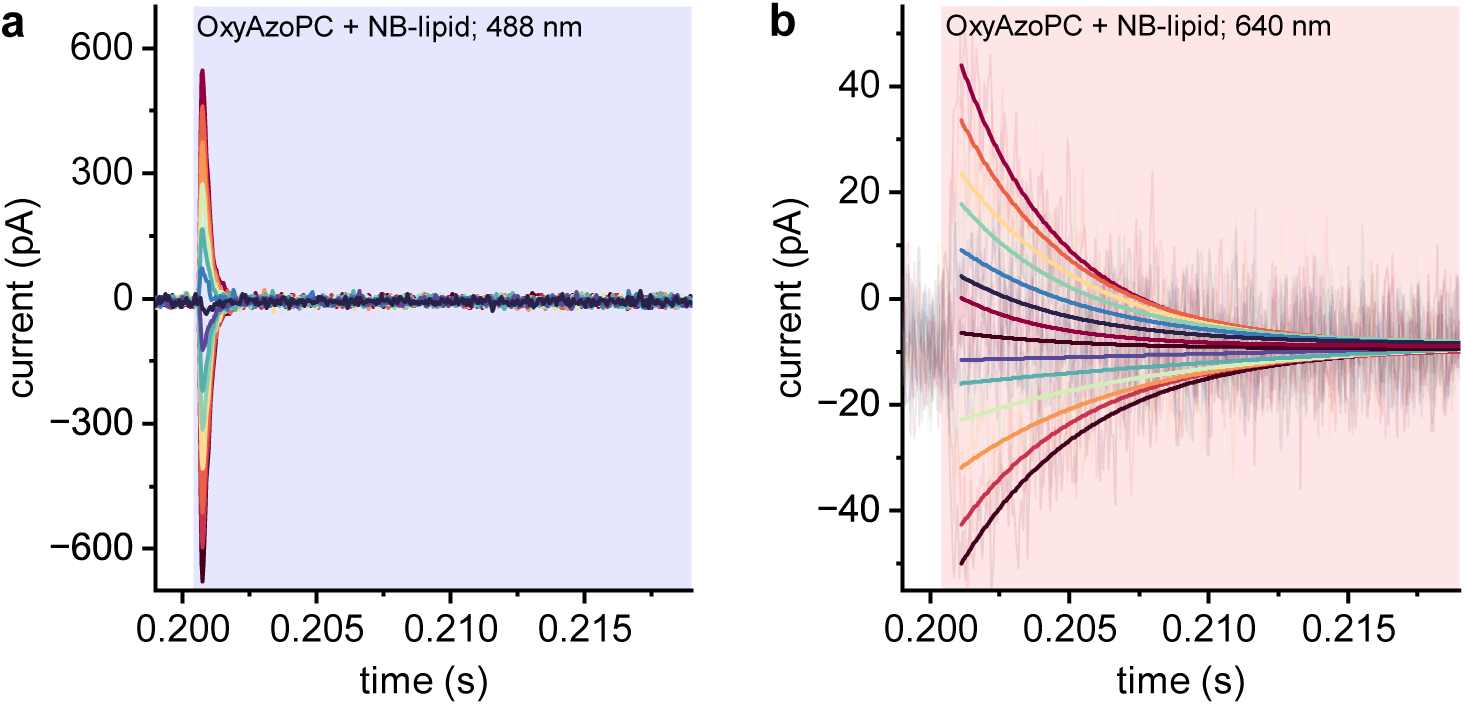
**a,** *E. coli* PLE bilayers containing 20 wt% *cis*-OxyAzoPC and 1 wt% NB-lipid in both leaflets exposed to blue light exhibit optocapacitive currents, with *τ* = 0.24 ms. The applied voltage was −130 mV to +130 mV with Δ*V* = 20 mV. **b,** Likewise, red-light illumination of the same bilayer evokes optocapacitive currents. Their smaller magnitude and slower decay (*τ* ∼ 5 ms) reflect a reduction in photoisomerization rate and *dC*/*dt*.

**Figure S4:**
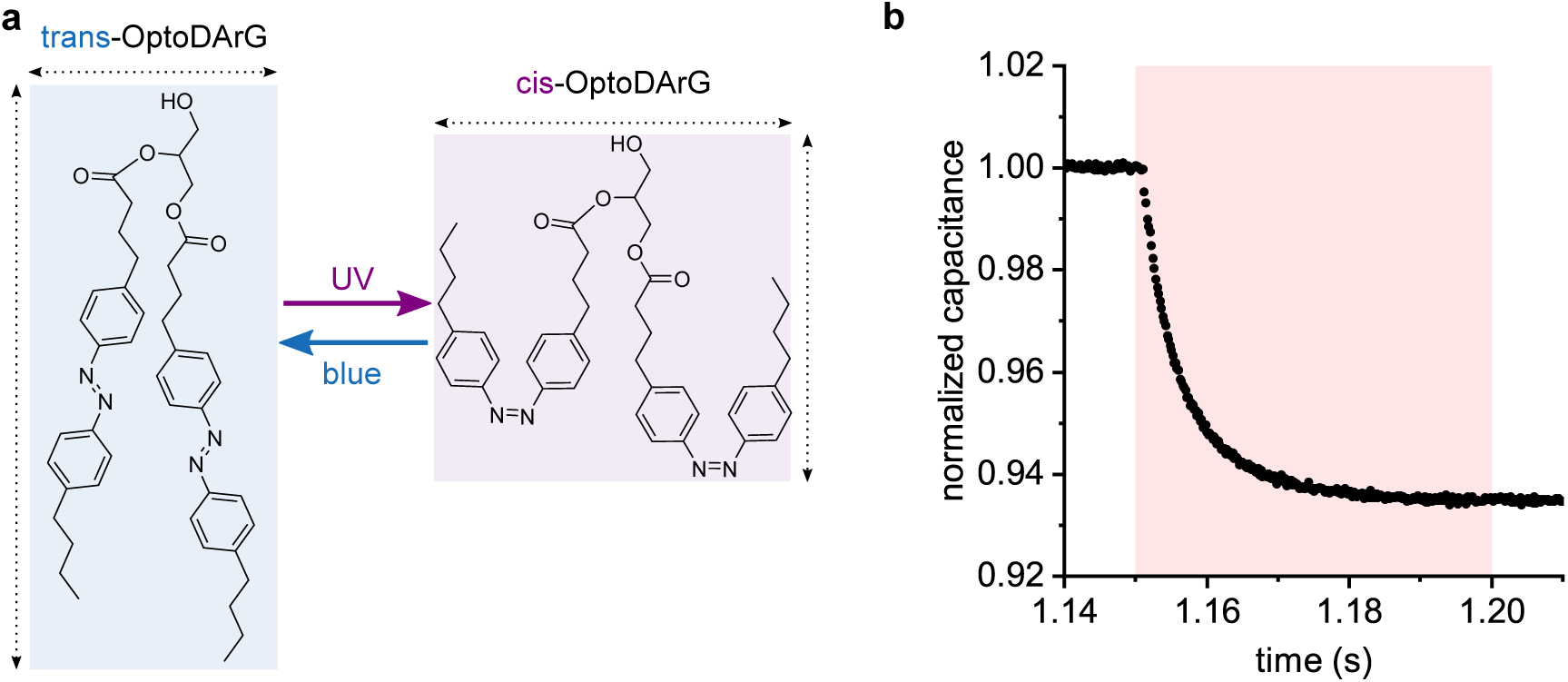
NB-lipid also catalyzes red-light *cis*→*trans* switching of other azobenzene photolipids in lipid bilayers. **a,** OptoDArG is a diacylglycerol-like photolipid with two *para,para*-dialkyl azobenzene moieties per molecule. It has previously been used for gating TRPC3 channels [42], modulating mechanosensitive peptide and ion channels [16, 21], modulating small-molecule carriers and membrane ion permeability [26], as well as eliciting depolarizing optocapacitive currents [20]. **b,** A symmetric bilayer containing 20 wt% *cis*-OptoDArG and 1 wt% NB-lipid in both leaflets was exposed to a 50 ms red-light pulse starting at 1.15 s. The pronounced capacitance response is consistent with OptoDArG photoisomerization.

**Figure S5:**
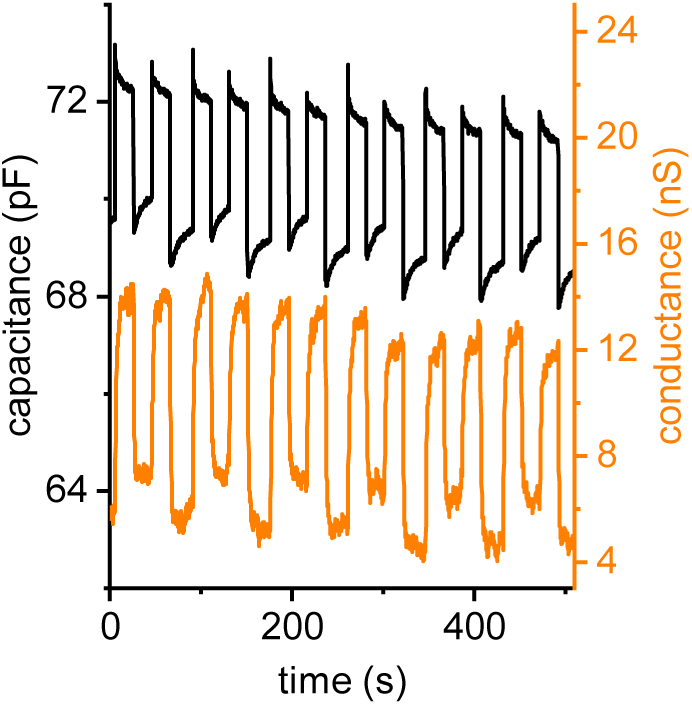
Repeated wavelength-selective switching does not show appreciable fatigue or loss of asymmetry. Full electrical record underlying the averaged traces shown in Figure 3c. The bilayer contained 20 wt% OxyAzoPC in both leaflets, 1 wt% NB-lipid in only one leaflet, and gramicidin A channels. Six consecutive UV-A/red/UV-A/blue sequences are shown, comprising 12 UV-A reset pulses and 12 red/blue actuation pulses. Each pulse lasted 50 ms. The difference between red-light-induced asymmetric and blue-light-induced symmetric switching remains preserved throughout the experiment without a progressive decrease in the capacitance response. The slow baseline drift in absolute capacitance and conductance occurs on a much longer timescale than the light-induced switching response and does not progressively alter the switching amplitudes. It is consistent with slow readjustments in PLB area which are usually observed; here, drift may be further enhanced by repeated light cycling which exerts push (UV-A) and pull (blue, red) on the surrounding lipid torus [21]. The concomitant conductance record likewise shows no abrupt or excessive cumulative loss of gA activity.

**Figure S6:**
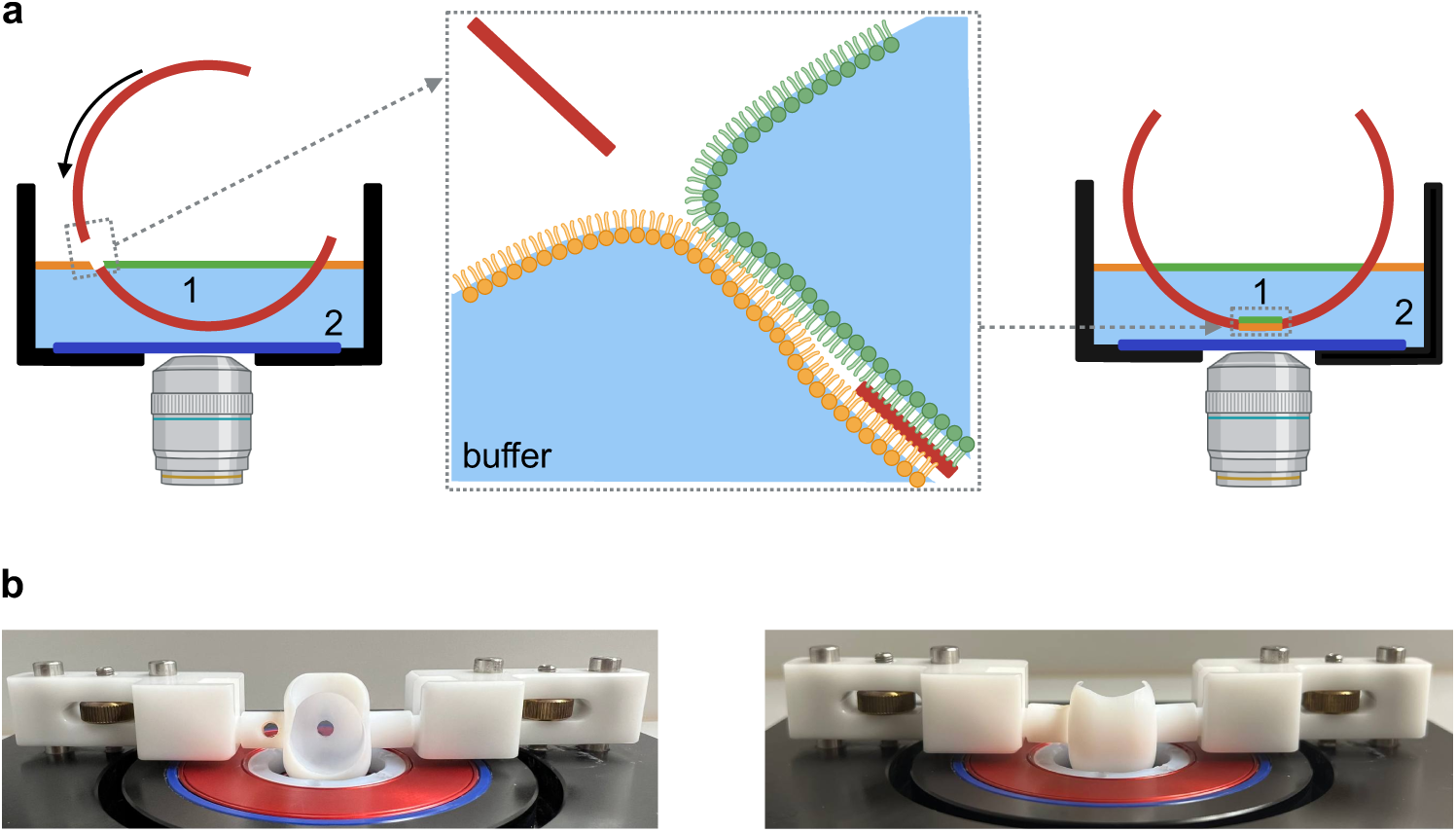
Schematic of asymmetric planar lipid bilayer formation by folding from distinct lipid monolayers. **a,** Different lipid mixtures (10 mg mL^−1^ in hexane) are first spread independently at the upper (compartment 1) and lower (compartment 2) air–water interfaces, creating a lipid monolayer in the upper compartment (shown in green) and a lipid monolayer in the lower compartment (shown in orange). Both compartments contain an aqueous buffer. After evaporation of the hexane, rotation of the PTFE diaphragm brings the two distinct lipid monolayers into apposition within the aperture. This action, called folding, brings together the two leaflets of the final asymmetric planar lipid bilayer, with the ‘green’ lipids on one side and the ‘orange’ lipids on the other. A residual hexadecane annulus surrounds and stabilizes the formed bilayer at the aperture rim. The upper compartment (red) is cylindrical and axially symmetric. The cross-section shown corresponds to a plane orthogonal to the cylinder’s axis. **b,** Below the schematic are photographs of the chamber for horizontal PLB formation.

